# Dynamics of methane cycling microbiome during methane flux hot moments from riparian buffer systems

**DOI:** 10.1101/2022.11.24.517857

**Authors:** Dasiel Obregon, Tolulope Mafa-Attoye, Megan Baskerville, Eduardo Mitter, Leandro Fonseca de Souza, Maren Oelbermann, Naresh Thevathasan, Siu Mui Tsai, Kari Dunfield

## Abstract

Riparian buffer systems (RBS) are a common agroforestry practice that consists of keeping a forested boundary adjacent to water bodies in agricultural landscapes, thus helping to protect aquatic ecosystems from adverse impacts. Nevertheless, despite the multiple benefits they provide, RBSs can be hotspots of methane emissions since abundant organic carbon and high-water tables are often found in these soils. In southern Ontario, Canada, the rehabilitation of Washington Creek’s streambank occurred in 1985. In a recent study, methane (CH_4_) fluxes were measured biweekly for two years (2017-2018) in four different vegetative riparian areas alongside Washington creek: a rehabilitated tree buffer (RH), a grassed buffer (GRB), an undisturbed deciduous forest (UNF), an undisturbed coniferous forest (CF), and an adjacent agricultural field (AGR) for comparison. Based on methane fluxes in 2018 and hot moments identified, we selected two dates from summer (July 04 and August 15) and use soil sampling from those days to assess the CH_4_ cycling microbial communities in these RBS. We used qPCR and high-throughput sequencing from both DNA and cDNA to measure the diversity and activity of the methanogen and methanotroph communities. Methanogens were abundant in all riparian soils, including the archaeal genera *Methanosaeta, Methanosarcina, Methanomassiliicoccus Methanoreggula*, but they were mostly active in UNF soils. Among methanotrophs, *Methylocystis* was the most abundant taxon in all the riparian sites, except for AGR soils where the methanotrophs community mostly comprised members of rice paddy clusters (RPCs and RPC-1) and upland soil clusters (TUSC and USCα). In summary, these results indicate that differences in CH_4_ fluxes between RBS at Washington creek are influenced by differences in the presence and activity of methanogens, which were higher in the deciduous forest (UNF) soils during hot moments CH_4_ flux, likely due to high water content in that soils.

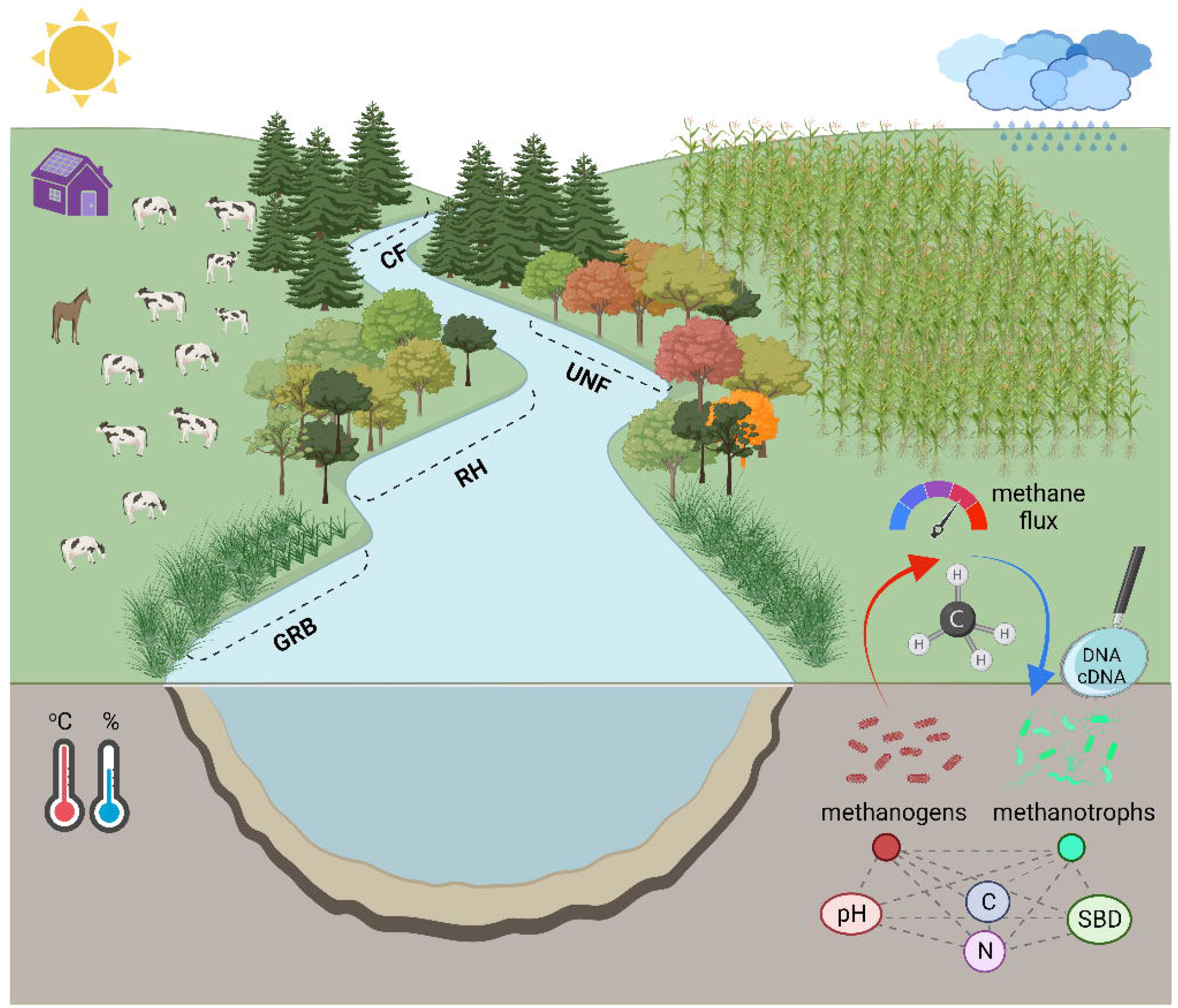

## 1 Introduction

Riparian buffers are the transitional boundary between terrestrial and aquatic ecosystems, usually forested, that help to protect the stream from the impact of adjacent land uses (NRCS, 2003). Freshwater ecosystems (e.g., streams, creeks and rivers) are among the most affected by the loss of riparian vegetation due to land-use change and agricultural intensification (Fortier et al., 2010; Meneses et al., 2015; Ye et al., 2014). The establishment of riparian buffer systems (RBS) has become a widely used strategy for protecting water bodies, with predominantly planted trees and shrubs along agriculturally degraded streams (AAFC, 2021; Guidotti et al., 2020; NRCS, 2011). Among other ecological services, RBS protects aquatic habitats by providing shade and maintaining water temperatures, providing detritus and woody debris for the soil biota, and trapping water runoff of pesticides, sediments, and fertilizers from adjacent areas (Bourgeois et al., 2016; Fortier et al., 2010; Lovell and Sullivan, 2006).

Due to their role in retaining nutrients, RBS are fundamental to biogeochemical cycles within the landscape (Fortier et al., 2010). It is believed that RBS can contribute to mitigating GHG emissions through large amounts of carbon (C) sequestration via plant and soil biomass (Audet et al., 2013; Capon et al., 2013). Yet, RBS can also be a source of GHG emissions through microbially mediated processes of C mineralization and methanogenesis (Vidon et al., 2016), whereas these processes are enhanced by favorable soil conditions in RBS (e.g., high levels of soil organic C (SOC) and high water tables are conducive to increasing CH_4_ emissions) (Audet et al., 2013; Bradley et al., 2011; Vidon et al., 2019). Accordingly, some studies had found that RBSs are responsible for a high proportion of CH_4_ release in agricultural landscapes (Dinsmore et al., 2009b; Jacinthe et al., 2015;

Vidon et al., 2015, 2014). The CH_4_ fluxes in the RBS are also affected by the type of vegetation present, with contrasting results about what vegetation type (e.g., herbaceous or treed RBS) better contribute to reduce CH_4_ emission (Dutaur and Verchot, 2007; Kim et al., 2010a). Therefore, it is important to determine the contribution of RBS to CH_4_ cycles to evaluate the trade-offs between economic and environmental benefits (Sonesson et al., 2020; Vidon et al., 2015, 2019).

Methane cycling in soils is driven by anaerobic methanogens (archaea), and aerobic methanotrophs, which are mostly methane oxidizing bacteria (MOB) (Knief, 2015; Malyan et al., 2016). Several soil factors, such as moisture, temperature, pH, organic matter (OM) content, and nutrient availability (i.e., C, N, Cu, Fe), are known to affect CH_4_ synthesis and oxidation (Serrano-Silva et al., 2014; Tate, 2015). Since groundwater level is a key driver of O_2_ diffusion, both methanogenesis (anaerobic) and methanotrophy (aerobic) are distinctly affected by soil water content (Malyan et al., 2016; Matson et al., 2017; Serrano-Silva et al., 2014). Consequently, several studies have reported negative correlations between water table depth and CH_4_ fluxes in RBS (Dinsmore et al., 2009a; Jacinthe et al., 2015). Yet, only a few studies have addressed the methanogens and methanotrophs communities in the context of riparian buffers (Kim et al., 2008; Krause et al., 2013).

Temperate riparian buffers are characterized by temperature variability and seasonal water table, in which their interaction are key drivers of CH_4_ flux (Jacinthe et al., 2015; Kaiser et al., 2018; Mander et al., 2015; Vidon et al., 2015). For instance, in RBS in the United States Midwest, Vidon et al. (2014) reported strong CH_4_ emissions after intense summer flooding events. These results were also confirmed by Jacinthe et al. (2015), who detected high CH_4_ emission rates after rewetting events in the summer (>22°C), but not after spring floods, likely due to low (<11°C) soil temperatures. Therefore, it is expected that spatial (‘hot spots’) and temporal (‘hot moments’) variability in CH_4_ fluxes occur between RBS in response to changes in edaphic factors (e.g., pH, C, SBD), but also changes in environmental conditions, either seasonally or throughout a weather event (Jacinthe et al., 2015; Vidon et al., 2015, 2014).

In Canada, one of the first experimental studies of riparian rehabilitation was established in 1985 adjacent to an agriculturally degraded stream (Washington Creek) in southern Ontario (Oelbermann and Gordon, 2000). Beneficial impacts of the RBS on both local terrestrial and aquatic environments were observed since rehabilitation was initiated, e.g., reduced soil respiration, enhanced wildlife habitat and maintenance of stream water temperature (Oelbermann et al., 2008). Concerning greenhouse gases, De Carlo et al. (2019) found no significant differences in N_2_O emissions between the rehabilitated and an upstream natural riparian forests, which was confirmed later by Baskerville et al. (2021), who also found lower N_2_O emissions in the RBS compared to an agricultural field adjacent to the rehabilitated RBS. Moreover, Mafa-Attoye et al. (2020) detected significant differences in N-cycling bacterial communities between RBS and agricultural soils. Recently, a study conducted by Baskerville et al. (2021) quantified the seasonal variation of CO_2_ and CH_4_ fluxes for two years (2017 and 2018) along Washington Creek. Specifically, they performed biweekly gas measurements and detected differences in CH_4_ fluxes among RBSs as well as seasonal variability within them, based on different types of vegetation.

For further understanding of the biogeochemical CH_4_ cycle in this context, and specifically the activity of the microbial communities driving the CH_4_ production and consumption, we identified hot moments of CH_4_ emission (representing CH_4_ flux peaks in the framework of this study) in the seasonal measurements in 2018 from Baskerville et al. (2021). Using the soil that was concurrently sampled during gas collection by Baskerville et al. (2021) we set out to characterize the methanogen and methanotroph communities in terms of abundance, taxonomic diversity, and activity. We hypothesize that different CH_4_-cycling taxa are ubiquitous in the different RBS soils and are driven by different edaphic factors and stimulated by environmental conditions (i.e., seasonal rainfall and temperature). This study aimed to *i*) evaluate methanogen and methanotroph abundance and community composition between RBS, and compare it to an adjacent agricultural field, *ii*) determine the taxonomic profile of active taxa during hot moments of CH_4_ flux, and *iii*) identify abiotic drivers of CH_4_-cycling communities.

## 2 Material and Methods

### 2.1 Study site

This study was conducted at Washington Creek, located in the Township of Blandford-Blenheim, Oxford County, Ontario, Canada. Washington Creek is a 9-km long 1st-order spring-fed stream within the Grand River watershed that flows into the Nith River south of Plattsville (43°18’N 80°33’W) (Figure 1). The landscape in Oxford County is dominated by agricultural fields with very little streambank vegetation, causing a high degree of streambank and aquatic habitat degradation. The climate is temperate, with a mean annual temperature of 7.3°C, mean annual precipitation is 919 mm (Supplementary Figure S1), and a mean annual frost-free period of 208 days (Environment Canada, 2020). The soil in Oxford County is classified as a Grey Brown Luvisol that has an overall loamy texture with hilly areas consisting of silt loam and sand, Plattsville is located 304 m above sea level (Mozuraitis and Hagarty, 1996). The agricultural landscape along Washington Creek is dominated by row cropping of corn (*Zea mays L*.) in rotation with soybeans (*Glycine max* (L.), or pastureland on both sides of the stream. For this study, four RBS with different vegetation types were sampled along Washington Creek: an undisturbed natural deciduous forest (UNF), a natural coniferous forest (CF), a grassed riparian buffer (GRB), a 33-year-old rehabilitated forest buffer (RH), and an agricultural (AGR) field on soybeans rotation (Figure 1).

**Figure. 1.**
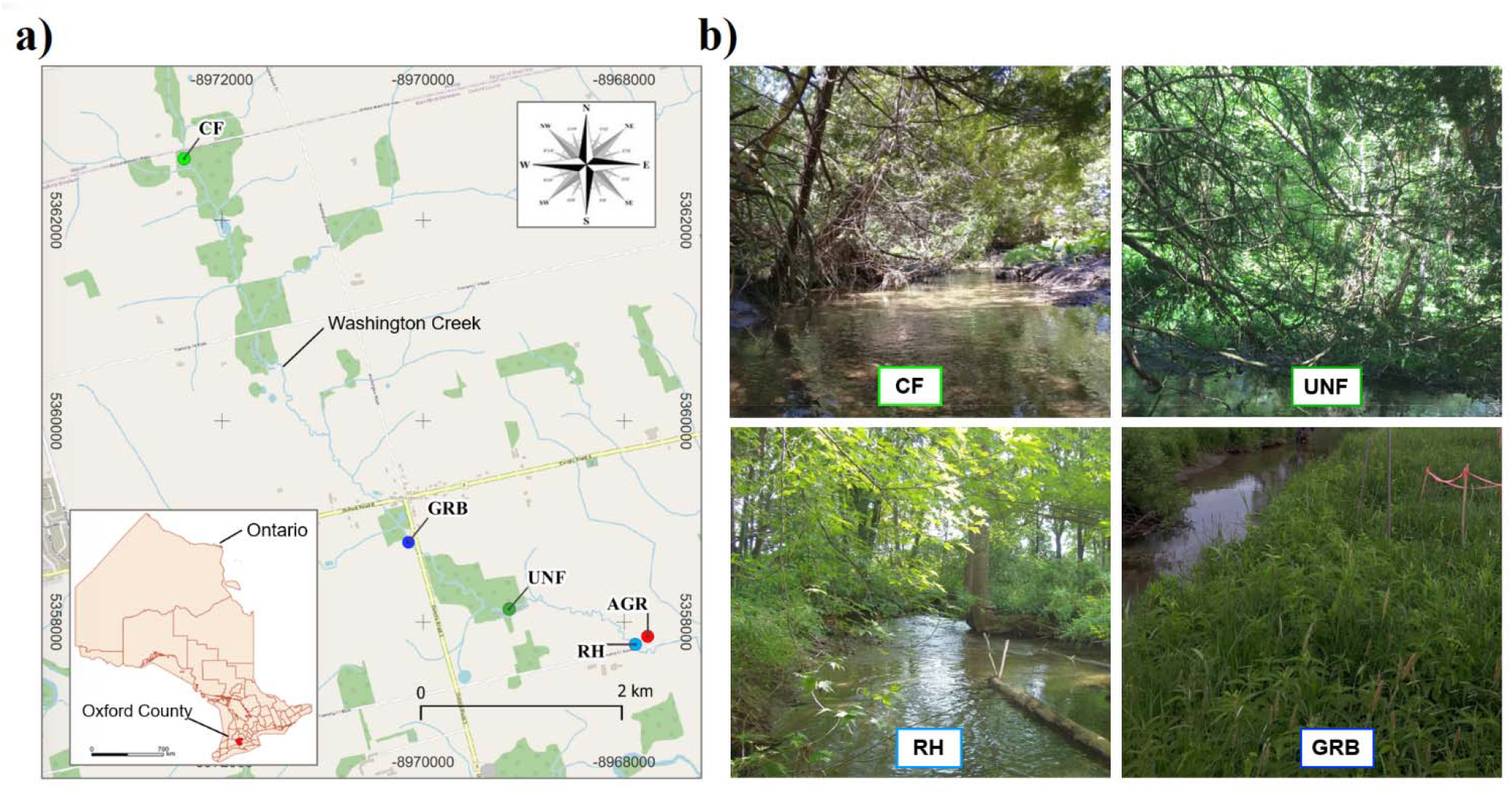
Sampling area. **a)** Aerial view of the Washington Creek region, located in the Township of Blandford-Blenheim, Oxford County, Ontario, Canada. Diverse RBS occur along the same agriculturally degraded stream. Four sites from different RBS were studied, which occur on both sides of the stream and are situated within a 5-km stretch of Washington Creek. The geographic location of each site is shown: CF-undisturbed coniferous forest, UNF-undisturbed deciduous forest, GRB-grassed buffer, RH-rehabilitated forest buffer, and AGR-agricultural land. **b)** A picture from each one of the RBS is shown, except for the AGR soil which consisted of a conventional soybean field.

Detailed information about the vegetation in each RBS site was provided by Baskerville et al. (2021). Briefly, the UNF RBS predominantly consisted of American beech (*Fagus grandifolia* E.), sugar maple (*Acer saccharum* L.), basswood (*Tilia americana* L.), and eastern hemlock (*Tsuga canadensis*) that has remained undisturbed for ∼150 years. The CF RBS occurs at the source of Washington Creek, and is composed of Eastern White Cedar (*Thuga occidentalis*). This RBS has been undisturbed for ∼100 years and is characterized by a high amount of deadwood with little understory vegetation. The RH RBS consisted of perennial vegetation such as alder [*Alnus incana* subsp. *Rugosa* (Du Roi) R.T. Clausen., *Alnus glutinosa* (L.) Gaertn., and *Alnus rubra* Bong.], hybrid poplar (*Populus* x *canadensis* Moench), silver maple (*Acer saccharinum* L.), multiflora rosevine (*Rosa multiflora* Thunb.) and red osier dogwood (*Cornus sericea* subsp. *sericea* L.) (Oelbermann et al., 2008). The grass buffer (GRB) consists mainly of bentgrass (*Agrostis app*.) and purple-stemmed aster (*Symphyotrichum puniceum*), this area separates Washington Creek from agricultural land. Finally, a conventional agricultural field (AGR) was included in the study for further comparison, the field is tile-drained under conventional tillage system, currently under a corn-soybean rotation.

### 2.2 Methane flux measurement and processing

Seasonal CH_4_ fluxes were measured at each site during the frost-free period (spring-summer-fall) from March to November in 2017 and 2018 (Baskerville et al. 2021). For this study, we focused on CH_4_ fluxes from March 8 to November 14, 2018. A detailed description of CH_4_ fluxes measurements procedures is provided by Baskerville et al. (2021), briefly, CH_4_ fluxes were measured biweekly in four static chambers that were randomly placed within a 5 × 30 m area within each RBS, directly adjacent to the stream’s edge. Static chamber anchors (25 cm height, 10 cm radius) were constructed of white PVC piping inserted into a 10 cm soil depth. Chamber caps were removed after each sampling time, and soil was exposed to air between sampling dates. Gas samples were analyzed using an Agilent 6890 Gas Chromatograph (Agilent Technologies, Inc., Santa Clara, CA, USA). CH_4_ flux (μg CH_4_ m^-1^ h^-1^) was determined by the linear or curvilinear equation model, according to the best fit (Hutchinson and Mosier, 1981). Cumulative CH_4_ emissions during the period (∼245 days) were estimated for purpose of better data visualization. Calculations included averaging CH_4_ flux between two consecutive sampling dates, multiplied by 24 hours plus the number of days elapsed betweeen that dates, and summing to the estimated values from previous sampling days.

### 2.3 Soil sampling

Soil samples were collected biweekly, concurrently with GHG flux measurements from March to November 2018. In each RBS four 1 m^-2^ sub-plots, adjacent to each GHG chamber, were established. Each time, four soil samples were collected from each sub-plot with a soil auger to a 10 cm depth. 2 g of soil was immediately transferred into pre-weighed sterile tubes containing 3 ml of LifeGuard soil preservation solution (Qiagen, Toronto, CA) to stabilize the RNA. Tubes were immediately stored on ice, transported to the laboratory, stored -80 °C and extracted for soil RNA and DNA within 3 weeks. The remaining soil was stored for physicochemical analyses at 4 °C. For this study, we selected two sample dates in 2018 (July 04 and August 15; hereafter referred to as “Jul04” and “Aug15) for soil physicochemical and microbial analyses. The dates were selected based on CH_4_ fluxes differences among the RBS (i.e., on Jul04 and Aug15, some RBS had CH_4_ emissions peaks while others showed maximum consumption).

### 2.4 Soil physicochemical analysis

Soil moisture (%) and temperature (°C) were obtained from each soil sample location using an HH2-WET Sensor (Delta T Devices, Cambridge, UK). Measurements were made bi-weekly at the time of gas measurements and were taken to a 10 cm depth. Soil pH was determined on 10 g soil diluted in deionized water, stirred, and allowed to sit for 1 hour, pH was measured using calibrated pH meter. Total soil carbon (TC) and total nitrogen (TN) were determined using air-dried soil samples sieved to 2 mm and analyzed on an elemental analyzer (CN 628, LECO Instruments, Canada), these data was gently provided by Dr. Kira A. Borden (Borden et al., 2021). Available ammonium (NH_3_) and nitrate (NO_3_) were determined on 5 g of air-dried soil, mixed with KCl for 15 minutes at 180 rpm using a reciprocating shaker, then filtered through Whatman 42 filter paper, and analyzed using a Shimadzu 1800 UV-Vis Spectrophotometer (Shimadzu Corp., Kyoto, Japan) at 640 nm to determine NH_3_, and 540 nm to determine NO_3_. Soil bulk density (SBD) was determined from collected soil cores with predetermined volume. The soil was weighed, dried at 105°C for 4 days, and reweighed after drying.

### 2.5 RNA and DNA extraction

Soil samples in Lifeguard preservation solution were centrifuged, and the sediments were used to coextract the total RNA and DNA using the RNeasy PowerSoil Total RNA isolation kit and the DNA elution kit according to the manufacturer’s protocol (Qiagen^®^, Valencia, CA). To eliminate potential contaminant DNA in RNA samples, RNase-free DNase (Promega GmbH, Mannheim, Germany) was added to 8μl of RNA in reaction tubes in triplicates. The purified RNA was reverse transcribed into single-stranded complementary DNA (cDNA) using the Applied Biosystems^®^ High-Capacity cDNA Reverse Transcription Kit (Life Technologies, Burlington, Canada) as recommended by the manufacturer. Both cDNA and DNA were quantified using a Qubit ™ 4.0 fluorometer (Life Technologies, Burlington, Canada) and samples were subjected to inhibitory tests to determine appropriate dilutions for quantitative real-time PCR (qPCR).

### 2.6 Quantitative PCR

The abundance of methanogens and methanotrophs was assessed through qPCR assays targeting the marker genes *mcr*A, and *pmo*A and *mmo*X, respectively, in both DNA and cDNA. For each assay, standard curves were constructed using ten-fold serial dilutions based on gBlocks gene fragments (Blazejewski et al., 2019) constructed for each target sequence (gBlocks™, Integrated DNA Technologies, Inc, USA) following MIQE guidelines for qPCR assays. The primer sets and thermocycling conditions are presented in Supplementary Table 1. The qPCR reaction consisted of 10 μL of 1× SYBR green supermix (Bio-Rad Laboratories, Inc.), 1 μL (10 μM) of each forward and reverse primers, 2 μL of DNA or cDNA template (1 to 10 ng/μL) and 6 μL of DNase-free water to make a final volume of 20 μL. The results were expressed in Log gene copy numbers per g of dry soil (gene copy g^-1^).

### 2.7 High-throughput amplicon sequencing

High-throughput sequencing of CH_4_-cycling communities was performed on both DNA and cDNA from soil samples collected on Jul04. In order to achieve high sequencing coverage of methanogens and methanotrophs, different sequencing strategies of amplicon sequencing were used for each group. For methanogens, libraries were constructed using PCR products obtained after amplification using the primer set Arch340F/806rb (Supplementary Table 1) that targets specific V4/V5 region of archaeal 16S rRNA gene, which allows for obtaining ∼75% of archaeal sequences (Bahram et al., 2019). For methanotrophs, libraries were constructed using the *pmo*A gene; to target the broad diversity of *pmo*A sequences, a nested PCR (nPCR) procedure was adopted as described by Deng et al. (2019). Briefly, the primer set A189-A682r (Supplementary Table 1) was used in the 1st round as they provide broader coverage of *pmo*A diversity, but these primers also detect *amo*A sequence (from ammonia-oxidizing bacteria). Then, a specific primer set was used in the 2nd round of nPCR, which included the forward primerA189f and two reverse primers mb661r/650r (Supplementary Table 1). Library preparation and sequencing were performed according to standard protocols at Génome Québec Innovation Centre from McGill University, Montréal (Québec) Canada. High-throughput sequencing was performed using an Illumina MiSeq platform (Illumina Inc., USA).

### 2.8 Bioinformatics and sequencing data processing

Demultiplexed reads packages were pre-processed and analyzed using the QIIME2 pipeline v. 2021.8 (Bolyen et al., 2019). Briefly, data cleaning and merging of mate reads, including chimeras removal, was performed using DADA2 via q2-dada2 (Callahan et al., 2016); this approach enables sequence analysis resolution down to the single-nucleotide level, thus resolving each amplicon sequence variant (ASV). To explore the phylogenetic diversity in each data set, the representative ASVs were aligned using the MAFFT algorithm (Katoh, 2002) via q2-alignment plugin, then the alignments were used to construct the phylogeny following FastTree2 method (Price et al., 2010) via q2-phylogeny. The phylogenetic trees were visualized and edited using the iToL platform (http://itol.embl.de) (Letunic and Bork, 2016).

Taxonomic assignment of ASVs was performed using QIIME2 q2-feature-classifier plugin (Bokulich et al., 2017), however different strategies were used for each marker. In archaeal 16S sequences, taxonomic identification was performed using the Classify-Sklearn Naive Bayes method, using a pre-trained classifier (99%) based on the primers Arch340F/806rb and 16S rRNA SILVA database v.138 (Quast et al., 2012). For *pmoA*, quality filtered ASVs were annotated (99%) following the classify consensus Vsearch method, which is based on the alignment of query sequences to a reference sequence panel. As a reference panel, we used a *pmo*A gene sequence database, available online at GFZ Data Services platform (Yang et al., 2016).

### 2.9 Statistical analyses

The experimental design in this study is pseudo-replicated since no other stream with similar riparian zone age, vegetation types, rehabilitation practices, land-use management, and environmental conditions existed in the region under study; we acknowledge that this limits the universality and applicability of our results. Statistical analyses on CH_4_ fluxes were performed by Baskerville et al. (2021). Briefly, Linear mixed models (LMMs) were run to determine differences in CH_4_ flux among RBS and seasons. The qPCR data were compared between land uses using *t*-test for multiple comparisons, including Holm-Sidak correction (alpha=0.05). Differences between soil parameters (e.g., TC, TN, C/N, SBD, and pH) as determined by days July 04 and August 15 were compared using Tukey’s multiple comparison test (alpha=0.05) and displayed using principal component analysis (PCA).

Microbial diversity of the methanotrophic and archaeal communities was compared between land uses, based on phylogenetic metrics of alpha (i.e., Faith Phylogenetic Index) and beta (i.e., unweighted UniFrac distance) diversity. Comparison of alpha and beta diversity metrics were performed using pairwise Kruskal-Wallis and PERMANOVA tests, respectively. Principal coordinates analyses (PCoA) based plots on unweighted UniFrac distances derived from DNA and cDNA sequencing data were compared using Procrustes analysis, as implemented in QIIME2. In addition, the correlations between unweighted UniFrac distance matrix and soil properties distance matrix were analyzed using Mantel’s test based on Spearman’s rank correlation coefficients.

Co-occurrence networks were used to analyze and visualize associations of CH_4_ flux, soil properties and soil microbes. Networks were generated using the CoNet alpha plugin (http://psbweb05.psb.ugent.be/conet) (Faust and Raes, 2016) in Cytoscape v.3.8.0 (Shannon, 2003). CoNet is an ensemble□based network inference tool designed to detect non-random patterns of microbial co□occurrence using multiple correlations and similarity measures. Briefly, networks were calculated on methanogen and methanotroph sequencing data (genus level) across 15 samples, filtered to a minimum of 5 and 10 reads for cDNA and DNA, respectively. An additional metadata with soil properties was added as a “feature matrix” in order to detect relationships between taxa and environmental variables. Pairwise associations among taxa/soil properties were calculated using Pearson, Spearman and Kendall correlation methods simultaneously (1000 permutations). Edges were retained when supported by at least two correlation methods (coefficient > 0.35) and *p*□values below 0.05. Networks visualized with Gephi v.0.9.2 (Bastian et al., 2009).

## 3 Results

### 3.1 Seasonal CH_4_ fluxes from different riparian buffer systems at Washington creek

Seasonal CH_4_ fluxes in 2018 ranged from -140 to 1233 μg CH_4_-C m_-2_ h^-1^. The average CH_4_ fluxes in the UNF site, with a mean of 1233 μg CH_4_-C m_-2_ h^-1^ were significantly higher (*p*<0.001) compared to GRB (−6 μg CH_4_-C m_-2_ h^-1^), RH (−61 μg CH_4_-C m_-2_ h^-1^), CF (29 μg CH_4_-C m_-2_ h^-1^), and AGR site (−120 μg CH_4_-C m_-2_ h^-1^). Accordingly, cumulative CH_4_ emissions were observed at the UNF site, whereas the CH_4_ flux was kept close to zero in the CF RBS soils, and a trend to CH_4_ uptake was observed in GRB, RH and AGR sites (Figure 2a). Furthermore, several hot moments of CH_4_ fluxes were detected at the UNF RBS in all the seasons, and less frequent hot moments of CH_4_ fluxes were observed at the CF RBS site, and even at GRB, RH and AGR soils (Figure 2b). Based on these data, we selected July 04 (Jul04) and August 15 (Aug15) sampling dates for microbial analyses, specifically because contrasting CH_4_ fluxes were recorded in some RBSs on those days (i.e., methane emission peaks at UNF and higher CH_4_ uptake rates at GRB).

**Figure 2.**
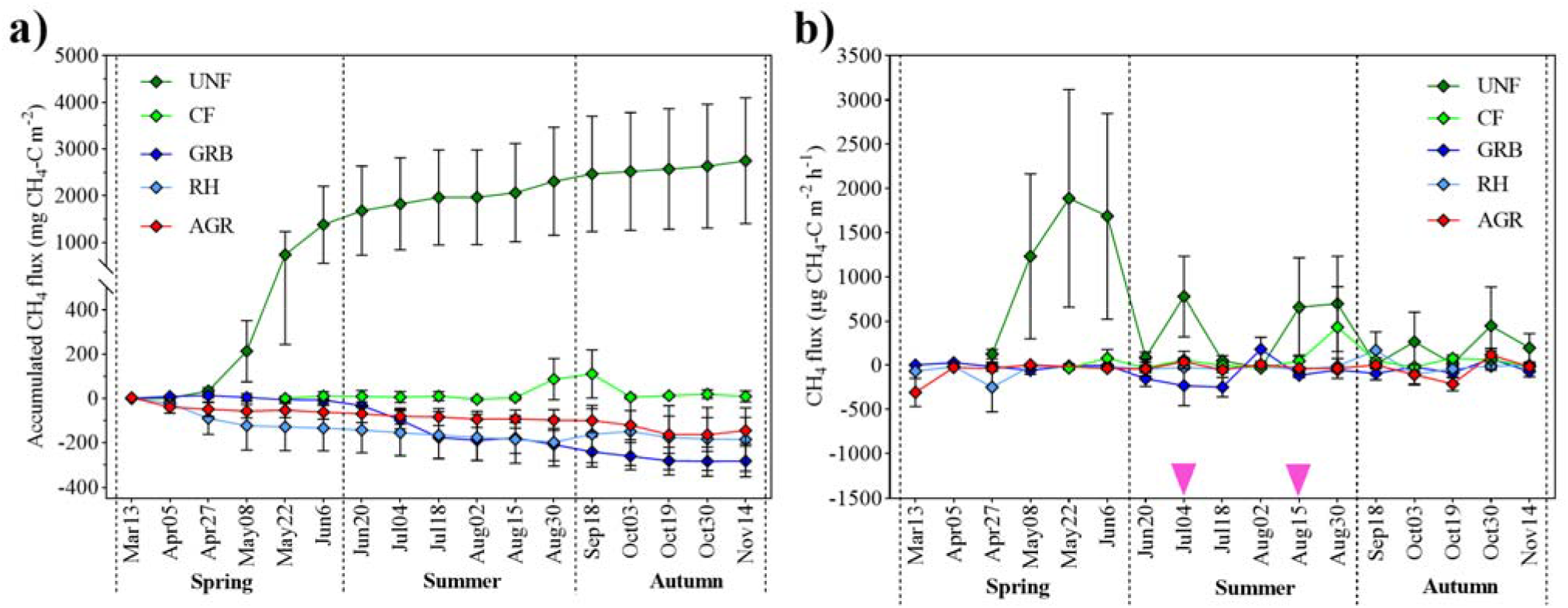
Seasonal CH_4_ flux during the frost-free period at different riparian buffer systems in the riparian zone of Washington creek in southern Ontario, including a grassed buffer (GRB), a rehabilitated forest buffer (RH), an undisturbed deciduous forest (UNF), a coniferous forest (CF), and adjacent agricultural field (AGR). Bi-weekly measurements of CH_4_ flux were conducted in four chambers at each site. **a)** Cumulative CH_4_ emissions corresponding to the full sampling period. **b)** Methane flux recorded on each sampling day. The dots represent the mean and the standard mean error. July 04 and August 15 were the sampling dates selected for soil microbial analyzes.

### 3.2 Abundance of methanogens and methanotrophs across different RBSs

Methanotroph abundance determined by *pmo*A and *mmo*X gene quantities was lowest in AGR soil compared to forests (UNF and CF) and rehabilitated tree buffer (RH) for both sampling days (Figure 3a,b). Specifically, methanotrophs harboring *pmo*A gene were less abundant in AGR than in GRB (Jul04: *p*=0.003; Aug15: *p*<0.001), and RH, CF, and UNF (*p*<0.001). Similar results were observed in *mmo*X gene abundance, except for less consistent differences between an AGR and GRB (Jul04: *p*=0.18; Aug15: *p=*0.02). Besides, the abundance of methanogens (*mcr*A gene) was higher in CF soils on Jul04. On Aug15 the abundance of *mcr*A was higher in forest CF and UNF soils. On the other hand, gene expression based on cDNA revealed a similar pattern among different land uses, with lower gene transcripts of *pmo*A and *mmo*X in AGR soils, and the highest abundance of *pmo*A and *mcr*A at CF and UNF sites (Figure 3c,d).

**Figure 3.**
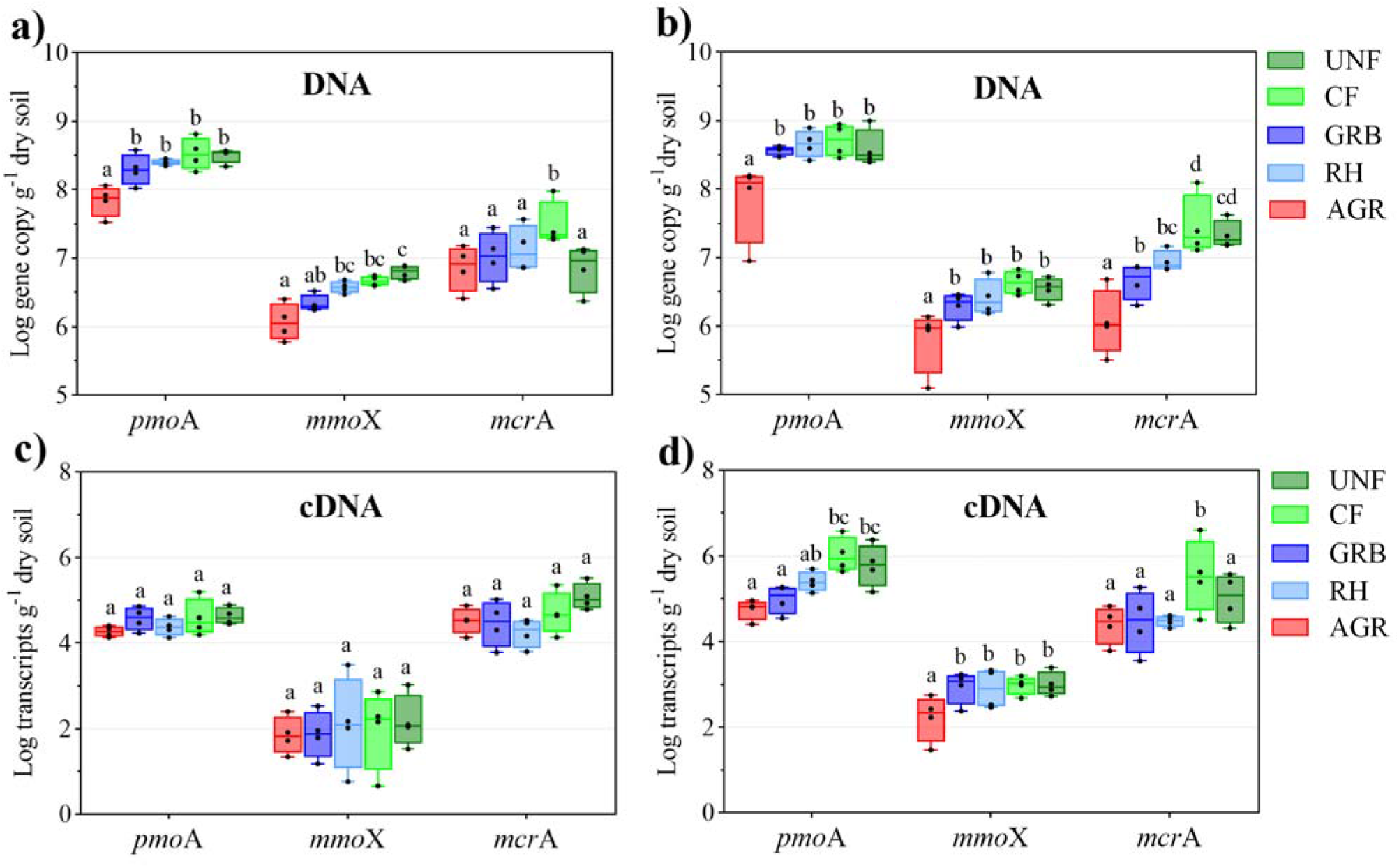
Abundance of the *mcr*A (methanogens), and *pmo*A and *mmo*X genes (methanotrophs) at different land-use sites. The quantification was performed using qPCR assays targeting DNA from soil samples collected on **(a)** July 04 (Jul04), and **(b)** August 15 (Aug15). For cDNA samples, quantifications of gene transcripts are presented for **(c)** Jul04 and **(d)** Aug15. Different letters within the same group (i.e., for each gene) indicate significant differences between land uses (Tukey HSD, *p*< 0.05)

### 3.3 Community composition and activity of methanogens in the different RBSs

A total of 411 and 325 ASVs were annotated as archaea in DNA and cDNA data, respectively. These ASVs belonged to at least 14 well-defined archaeal orders, in which the order Woesearchaeales comprised a large proportion of ASVs in both datasets (Supplementary Figure S2). Total archaeal community composition in AGR soils was significantly different from those in the forest (CF and UNF) and riparian (GRB and RH) soils in both DNA (*F*=1.82; *p*=0.002) and cDNA (*F*=1.45; *p*=0.04) (Supplementary Figure S3). Overall, archaea from the order Nitrososphaerales comprised most of the community profile in both DNA and cDNA data (Supplementary Figure S4). Besides, Nitrososphaerales, Woesearchaeales and Nitrosopumilales were also highly abundant in DNA sequencing data, whereas cDNA profiles had a high abundance of Bathyarchaeia and the methanogenic order Methanosarciniales. In further analysis, we selected the ASVs that were annotated as methanogens.

Methanogens from six orders were detected in both DNA and cDNA datasets (Figure 4). A total of 33 ASVs in the DNA data were classified as methanogens, among which the most abundant were the genera *Methanosaeta* and *Methanosarcina* from the order Methanosarciniales (36%,12/33), followed by 8 ASVs (24.2%) as *Methanomassiliicoccus* (Methanomassiliicoccales), and 7 (21.2%) as *Methanoregula* (Methanomicrobiales). Overall, the highest abundance of that ASVs was detected in UNF soils (Figure 4a). In cDNA data, a higher number of ASVs (total of 52) were assigned as methanogens compared with the DNA data set (Figure 4b), wherein Methanosarciniales were also the most abundant taxa (36.5%, 19/52). A higher number of ASVs from the order Methanomicrobiales (38.5%, 20/52) was observed also in the cDNA dataset, which most often consisted of the genera *Methanoregula* and *Methanospirillum*. Consistent with the distribution of the methanogens across land use from DNA data, the highest abundance of active methanogens (63.5%) was found at the UNF soils, thus consistent with the highest methane emission detected at this site.

**Figure 4.**
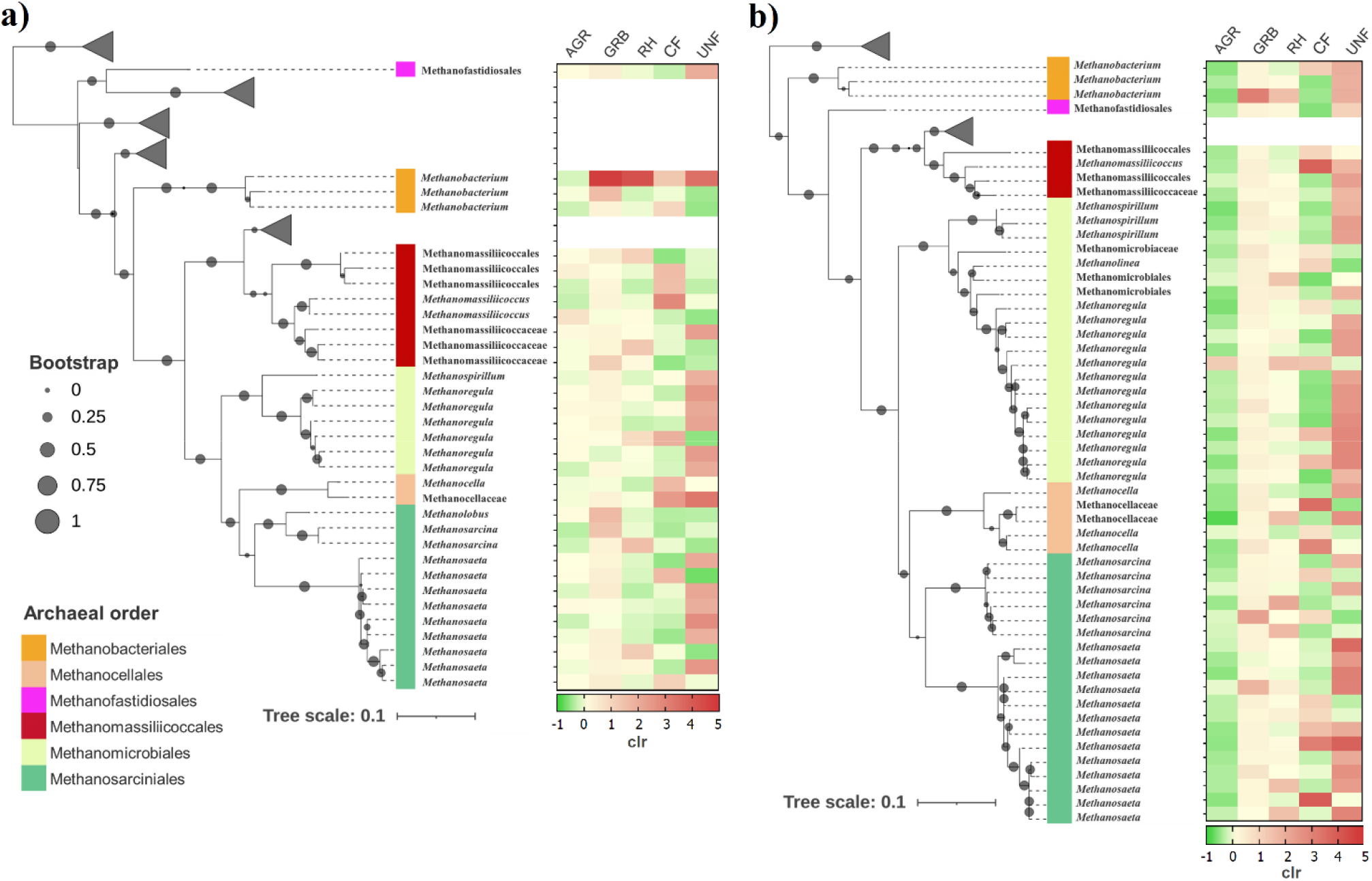
Maximum likelihood phylogenetic tree of ASVs from methanogens detected by DNA and cDNA sequencing from different land uses. Phylogenetic tree from **(a)** DNA and **(b)** cDNA sequences assigned as methanogens. The ASVs are identified at the order level and classified as genus or family when possible. Other clades containing non-methanogenic archaea are collapsed. In the heatmaps, the ASVs are presented as the average of their proportional distribution (clr: centered log-ratio transformed) among samples in each land use.

### 3.4 Diversity and taxonomic composition of the methanotrophic community

A difference in taxonomic richness of the *pmo*A DNA was observed between AGR and GRB soils (*H*=4.1; *p*=0.04). However, no significant differences were also observed between land uses in cDNA-based *pmo*A sequencing. No significant differences were found in community evenness between soils from both DNA and cDNA data (Supplementary Figure S5). Analysis of ß-diversity (i.e., based on unweighted UniFrac distances) revealed significant differences between AGR and riparian soils as confirmed on both DNA (*F*=3.45; *p*=0.001) and cDNA (*F*=2.26; *p*=0.02) sequencing data (Supplementary Figure S5).

Distinct taxonomic composition of the methanotroph community was observed in AGR soils compared to riparian (UNF, CF, GRB and RH) soils. On DNA-based sequences, a total of 297 ASVs were detected, which were classified into 13 taxons (at genus level) mostly from the phylum Proteobacteria (99%, 294/297), split into the classes Alphaproteobacteria (72.7%, 216/297) and Gammaproteobacteria (26.3%, 78/297). In addition, 1% (3/297) of the sequences were identified as “environmental_samples” at phylum level, and as the class “uncultured_bacterium (pxmA)”, thus identified as unclassified_Bacteria (MOB_like) for purposes of this work. At genus level, most ASVs (66.6%, 198/297) were identified as *Methylocystis*, followed by Tropical Upland Soil Cluster (TUSC_ pxmA) (9%, 27/297), type Ib methanotrophs belonging to Rice Paddy Cluster 1 (RPC-1) (7%, 21/297), type Ib (FWs) (3%, 9/297), typeIb (RPCs) (2.6%, 8/297) and type Ib Lake Washington Cluster (LWs) (2.3%, 7/297) (Figure 5a). Comparing land use, *Methylocystis* was mostly detected (> 98%) in riparian zones (UNF, CF, GRB and RH), whereas AGR soils had a high proportion of TUSC (pxmA), type Ib rice paddy cluster 1(RPC-1), type Ib (RPCs), followed by type IIb (USC-alpha) (Figure 5b).

**Figure 5.**
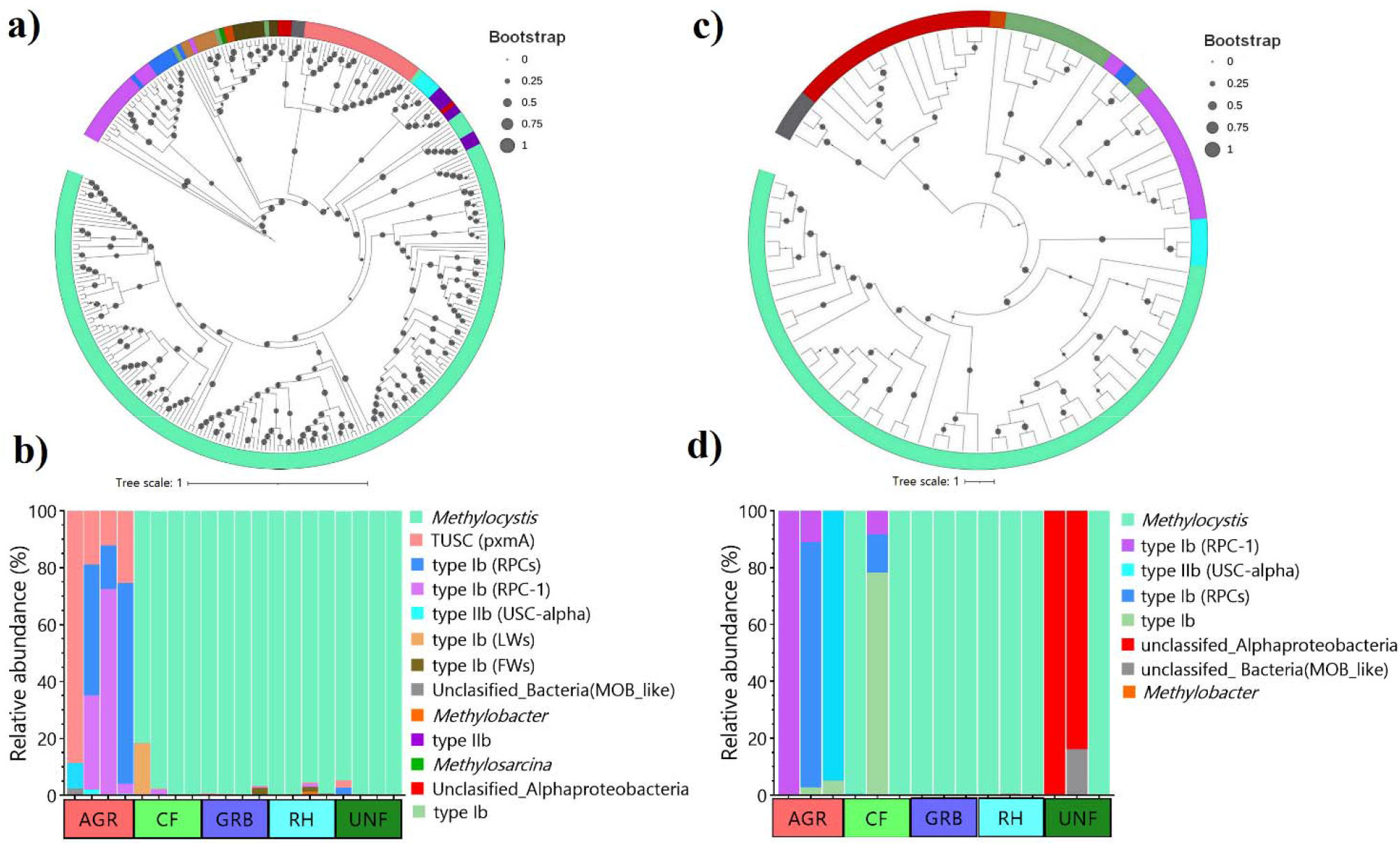
Taxonomic composition of methanotrophs at different land-use sites. The upper panel indicates the phylogenetic arrangement of *pmo*A sequences detected from **(a)** DNA and **(c)** cDNA sequencing data. The bottom panel shows the taxonomic composition at genus level based on both **(b)** DNA and **(d)** cDNA from each sample across five land use sites.

With cDNA-based sequencing, due to low sequencing coverage, some replicates were lost during the sequence denoising process. A total of 86 ASVs were detected, from which 74.4% (64/86) and 23.2% (20/86) were Alphaproteobacteria and Gammaproteobacteria respectively, while 3.5% (3/87) were annotated as unclasified_Bacteria (MOB_like). Similar to DNA profiles, *Methylocystis* comprised most of the ASVs detected at genus level (54.7%, 47/86), followed by unclasified_Alphaproteobacteria (15.1%, 13/86) and Type Ib (RPC-1) (10.4%, 9/86) (Figure 6c). The taxonomic composition of the methanotrophs in AGR soils was also distinct compared to all other riparian buffer sites. *Methylocystis* was the predominant active methanotrophs taxa in CF, GRB and RH soils, while in AGR soil the most abundant were type Ib (RPC-1), Type Ib (RPCs) and type IIb (USCα). However, differently from DNA-based sequencing, the active methanotrophs in the UNF forest predominantly consisted of unclassified_ Alphaproteobacteria, and one sample comprised ASVs assigned as unclassified_Bacteria (MOB_like) (Figure 5d). These two taxa are likely to be closely related genetically as they consistently clustered together in phylogenetic trees from both DNA and cDNA sequences (Figure 5c). Yet, it was not possible to refine the taxonomic classification of these taxa to other hierarchical levels (i.e., order, family or genus).

**Figure 6.**
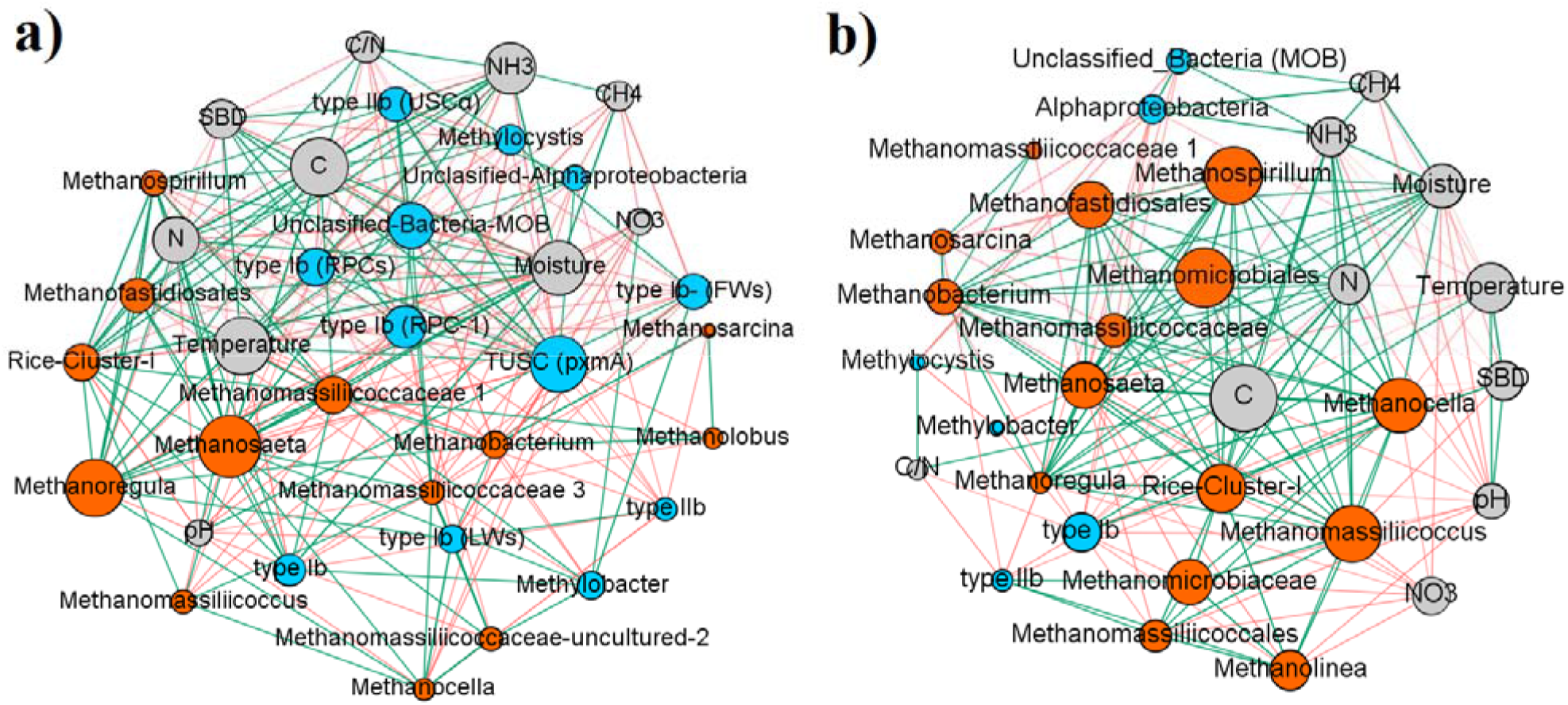
Correlation-based networks of methanogen and methanotroph communities and their interaction with soil properties. Networks were constructed from **(a)** DNA and **(b)** cDNA sequencing data. Node colors represent soil parameters (gray), methanogens (orange), and methanotrophs (blue), and node size reflects eigencentrality, which is indicative of the influence of the node on the networks. Edges between nodes represent significant negative (red) and positive (green) correlations, calculated from Pearson, Spearman, and Kendall correlation methods simultaneously (p< 0.05).

### 3.5 Analysis of edaphic factors across land uses

Analysis of soil temperature of the soils collected for microbial analysis revealed no significant differences between land uses (Table 1). Although not significant, the average temperatures detected in GRB and AGR soils were ∼2°C higher than in CF and UNF forests and RH soils. However, soil moisture was higher on sites with canopy coverage (UNF, CF and RH) when compared to other soils, especially in the UNF where soil moisture was three times higher than in GRB or AGR. Differences were also observed in TC in both sampling days, with a gradient from low TC content in AGR soils to high TC content in the CF and UNF. A similar gradient was observed for TN and NH_3_ from low content in AGR to higher contents forest sites (CF and UNF), although the differences were statistically significant only for NH_3_. Interestingly, the NO_3_ content was increased in GRB and RH but just on August 15. No significant differences were observed in soil C/N ratio or pH (*p*>0.05).

**Table 1.**
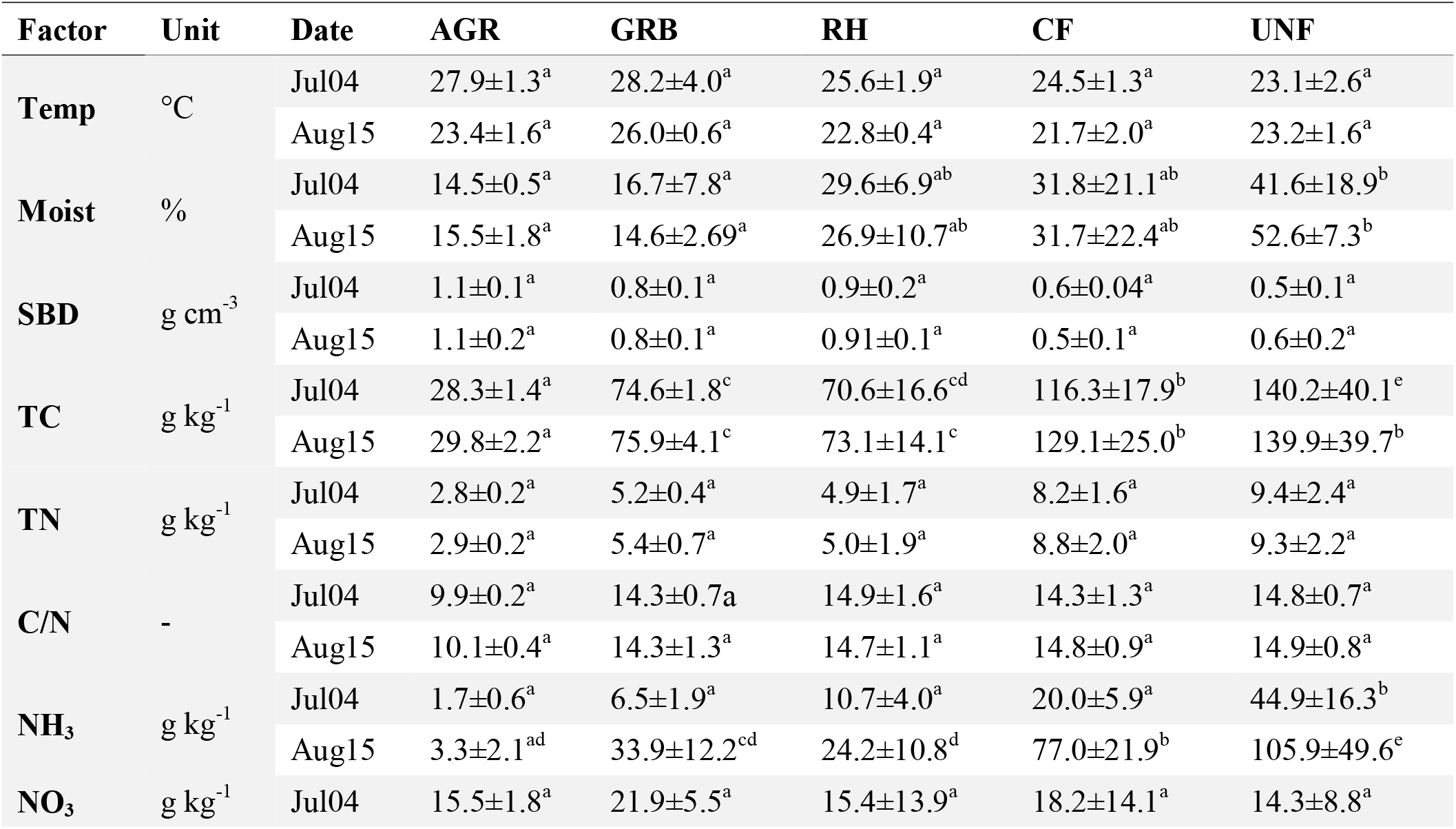
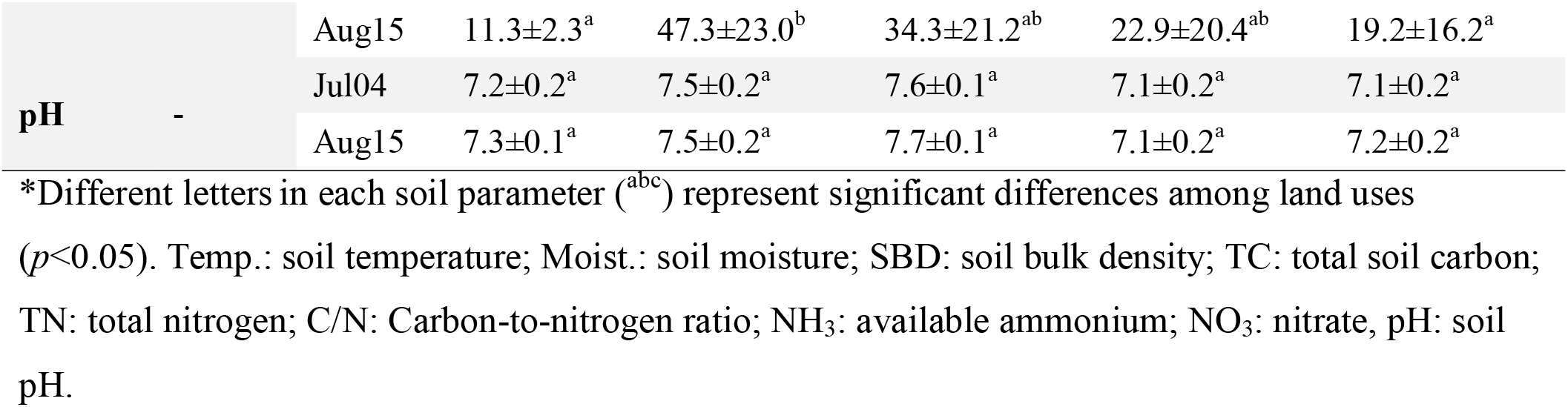
Soil properties from each RBS on average (mean and SD) over the soil sampling dates (July 04 and August 15).

Principal component analysis (PCA) based on soil edaphic factors, including CH_4_ flux, revealed significant differences (*F*= 6.30; *p*<0.001) between land uses. Overall, an overlap of riparian buffer soils (GRB and RH) was observed, associated with soil pH and temperature, whereas natural forest soils (UNF and CF) clustered together, associated with soil moisture and nutrient content. Agricultural soil samples showed a clear distance from other sites which was associated with SBD (Supplementary Figure S6).

### 3.6 Co-occurrence of methane cycling taxa and their association with soil edaphic factors

To explore the association between methane cycling taxa and soil conditions, we constructed co-occurrence networks based on three data matrices, including all the samples from all land uses: i) abundance of methanogens, ii) abundance of methanotrophs, and iii) soil edaphic factors, and the same analysis was performed for both DNA and cDNA sequencing data. Different patterns of microbe-microbe interactions, as well as between taxa and soil factors, were observed when DNA or cDNA sequencing data were explored (Figure 6). In the DNA-based network, soil C, temperature, moisture and NH_3_, were the most influential environmental factors, with eigencentrality of 0.91, 0.90, 0.85 and 0.78 respectively. Soil temperature had mainly positive correlations with methanotrophs [e.g., TUSC and type Ib (RCP-1)], while soil moisture was positively linked to methanogens (Figure 6a). Thirteen methanogens were detected in the DNA-based co-occurrence network, with an average degree (i.e., number o edges) of 15.2±6.7, whereas 12 methanotrophs were detected, with an average degree of 16.9±5.9 (Supplementary Table 2).

Instead of that, the methanogens were the most representative in cDNA-based networks, with a total of 15 nodes and an average degree of 13.7±5.3, while methanotrophs only constituted six nodes, with an average degree of 8.0±4.8 (Figure 6b). Soil C, temperature, and moisture, with eigencentrality of 1, 0.66, and 0.56 respectively, continues to have a strong influence within cDNA-based network, with positive correlations with most methanogens (Figure 6b; Supplementary Table 2). *Methylocystis*, although the most abundant methanotroph in these datasets, was not found associated with other taxa as shown in both DNA- and cDNA-based networks., in fact, *Methylocystis* generally exhibited negative correlations with other taxa, whereas it was positively linked to C/N ratio in both networks.

Interestingly, CH_4_ flux was correlated with soil moisture and NH_3_ in both networks, but it showed few associations with methanogens of methanotrophs.

## 4 Discussion

Riparian buffers are one of the most common agroforestry practices in Canada, and they are promoted to stabilize eroding banks of water bodies, but also to reduce nutrient runoff from adjacent agricultural land use, and protect the aquatic ecosystems (Oelbermann et al., 2008). However, RBS can also be source of CH_4_ emissions due to favorable environmental conditions (i.e., the presence of a high water table, and carbon addition as litterfall) (Dinsmore et al., 2009b; Jacinthe et al., 2015; Vidon et al., 2015, 2014). As previously described by Baskerville et al. (2021), amongst the RBS present at Washington Creek, southern Ontario, only the UNF RBS site acted as a hot spot of CH_4_ emissions, while all other RBS typically acted as CH_4_ sinks. Based on biweekly measurement of CH_4_ fluxes performed by Baskerville et al. (2021) during the frost free period in 2018, we identified hot moments of CH_4_ flux and selected two dates from summer (July 04 and August 15) for microbial analyses. Soil sampling from those days were used to assess the CH_4_ cycling microbial communities across these RBS. Specifically, we aimed to characterize the soil microbial communities involved in the production and consumption of CH_4_ in these RBS.

To further address the CH_4_-cycling communities, in this study we not only determined differences in abundance, taxonomic and phylogenetic composition of methanogens and methanotrophs but also considered the activity of these taxa during CH_4_ hot moments (i.e., high CH_4_ emission peaks in UNF versus high oxidation rates in GRB in the sampling dates under study). This approach is particularly important for methanogenic archaea (e.g., the genera *Methanosarcina* and *Methanocella*), which are ubiquitous in many upland soils worldwide, often in the dormancy stage but could be readily activated when anoxic conditions occur (Angel et al., 2012). Moreover, studies in the humid tropics suggest that soil biogeochemical dynamics can be accompanied by rapid shifts in CH_4_ fluxes (Fernandes et al., 2002; O’Connell et al., 2018; Verchot et al., 2000). For example, O’Connell et al. (2018) found that high CH_4_ uptake during drought periods was followed by a post-drought dramatic increase in CH_4_ emissions that offsets sink in a few weeks.

The abundance of methanogens was higher in deciduous forest UNF, with lower abundance in agricultural soils (AGR), as measured by qPCR from DNA and cDNA (Figure 3). Methanogens from six different orders were detected in most soils, except for agricultural soils, where other non-methanogenic archaeal orders (e.g., Nitrososphaerales) were predominant. We detected the presence of acetoclastic methanogens (i.e., *Methanosaeta* and *Methanosarcina*), but also hydrogenotrophic (i.e., *Methanoreggula* and *Methanomicrobium*) (Nazaries et al., 2013) and H_2_-dependent methylotrophic (*Methanomassiliicocus* and *Methanofastidiosa*) by DNA sequencing in all the RBS, mainly in the grassed buffer (GRB) and evergreen forest (CF). However, analysis of both DNA and cDNA sequencing data indicated that these taxa were most active in the UNF soil. These results not only coincide with the hot moments of CH_4_ emissions observed in these soils but also with the higth water content. The association between methanogens and soil moisture was also confirmed in co-occurrence networks, for example, an increased number of nodes from methanogens was detected in cDNA networks, which mostly were positively correlated with soil moisture, thus confirming the resilience of the methanogen community, which became active in response to favorable soil conditions.

To target the methanotroph community, we sequenced the functional gene *pmo*A instead of 16S rRNA gene, given that amplicon sequencing of the 16S rRNA gene often provides low sequencing coverage of methanotrophs since they constitute a small proportion within the total soil microbiota, and often it does not allow the identification of MOB taxa beyond known families (Knief, 2015). The *pmo*A gene is evolutionarily highly conserved and thus useful as a phylogenetic gene marker for methanotrophs (Holmes et al., 1995). In addition, phylogenetic trees constructed based on *pmo*A sequences closely reflect those of 16S rDNA based phylogenies for the same MOB organisms (Dunfield et al., 2002; Knief, 2015). Our results suggest that the genus *Methylocystis* widely predominates vegetative riparian buffers in Washington Creek, including natural forests. *Methylocystis* is considered to be a generalist MOB, inhabiting upland forest and grassland soils, even some *Methylocystis* species inhabit frozen soils and wetlands (e.g., *M. sporium* in arctic wetland soil and *M. sporium* in rice paddy soil) (Knief, 2015; Zeng et al., 2019). Several *Methylocystis* spp. harbour a paralog of the *pmo*A gene (i.e., *pmo*A2), which encodes the methane monooxygenase enabling methane oxidation at lower mixing ratios (i.e., high-affinity methane oxidation, HAMO) (Knief, 2015; Kolb, 2009). These capabilities provide resilience to this genera in hydromorphic soils where they often face low methane supply (Knief, 2015).

However, we did not detect *Methylocystis* in agricultural (AGR) soils. We assumed that the distinction of MOB community on AGR soils was most likely a consequence of land use, since frequent disturbances due to agricultural practices and land-use change are known to affect CH_4_ oxidation regardless of different climate and soil types (Tate, 2015). In particular, the non-detection of *Methylocystis* in AGR soils could be a consequence of changes in the C/N ratio, as our results revealed that only this taxon had positive correlations with C/N ratio in DNA- and cDNA-based networks.

Moreover, due to the competitive inhibition of methane monooxygenase by ammonia, changes in nitrogen (N) cycling are recognized to have strong effects on CH_4_ oxidation, thus affecting the abundance and diversity of methanotrophs (Nazaries et al., 2013; Tate, 2015). Yet, the dynamics between nitrogen and methane oxidation are not fully understood, as studies suggest that soil N, especially in the form of NH_4_^+^, could either stimulate, inhibit, or exert no influence on soil CH_4_ (Bodelier, 2011; Szafranek-Nakonieczna et al., 2019). In this study, type II Alpha-MOB such as *Methylocystis* ssp. thrive better in RBS soils with high NH_3_ and TN (e.g., CF and UNF).

These results are in agreement with (Nyerges et al., 2010) who found that *Methylocystis* spp. were more tolerant to the inhibitory effects of ammonium than nitrate. On the other hand, MOB in agricultural soils comprised mostly of type Rice Paddy Clusters (RPC-1and other RPCs) and Upland Soil Clusters (TUSC and USCα). The RPC clusters are usually detected in rice paddy-associated habitats, but they could also be heterogeneous and inhabit diverse environments (e.g., the large RPC1_3 cluster) (Knief, 2015). Surprisingly, TUSC and USCα in our study were more abundant in AGR soils, whereas these taxonomic groups are mostly associated with upland forest soils (Deng et al., 2019; Feng et al., 2020; Kou et al., 2020) and less frequent in intensively managed agricultural soils (Knief, 2015). Nevertheless, USCs have also been found in agricultural soils with high C content (Lima et al., 2014). Further studies are needed to address the drivers of the methanotrophs community at RBS at Washington Creek, Ontario.

The active methanotrophs in UNF RBS, were mostly only able to be classified at phylum level [i.e., unclassified_Bacteria (MOB_like) and unclassified_Alphaproteobacteria]; however, their presence confirms that methanotrophic activity persists in these soils even at high soil moisture. In addition, these taxa were the only nodes of methanotrophs that positively correlated with CH_4_ flux in the cDNA sequencing co-occurrence network. Our results are in agreement with Cai et al. (2016), who found that methane uptake is mediated by conventional methanotrophs even in periodically drained ecosystems. Yet, the fact that these taxa responded to increased soil moisture and high CH_4_ concentrations, but were taxonomically unidentified according to the reference database, may suggest a novel taxonomic group in these soils. Further studies using a more comprehensive sequencing approach (i.e., metagenomic sequencing) are needed in order to characterize the taxonomic, genomic, and functional traits of the methanotrophs in these ecosystems.

## 5 Conclusions

Riparian Buffer Systems (RBS) at Washington Creek, Southern Ontario, including rehabilitated forest buffers (RH), grassed buffers (GRB) and natural coniferous forest buffers (CF) have been proven to be sinks of atmospheric CH_4_, except for the natural deciduous forest buffer (UNF), which represents a hotspot for CH4 emissions, most of which occurred during spring and summer days, associated with high soil C content and high soil moisture levels. We confirm the hypothesis that different metahnogen and methanotroph taxa are ubiquitous in these RBS soils and are driven by different edaphic factors and environmental conditions. Methanogens from six taxonomic orders were present in all the RBS, with the highest abundance and taxonomic diversity at UNF, and associated with soil moisture and soil C content. Methanotrophs were also detected, and the main differences in taxonomic diversity were observed between RBS and the adjacent crop field (AGR). The *Methylocistys* spp. was discovered to be the most frequent methanotroph in RBS soils, but not in AGR soils where methanotrophs had low abundance and comprised mostly of members of Rice Paddy Clusters (RPC) and Upland Soil Cluster (USC). The findings of our study demonstrate the reliability of targeting both DNA and cDNA to measure diversity as well as the activity of CH_4_-cycling taxa for better understanding CH_4_ flux patterns. Overall, these results should be considered when establishing riparian buffer systems in agricultural landscapes, which should be designed not only based on the adjacent land uses and topographic conditions, but also on vegetative systems that help to diminish the activity of methanogens and thus reduce CH_4_ emissions.

## CRediT author contribution statement

**Dasiel Obregon:** Conceptualization, Methodology, Software, Formal analysis, Investigation, visualization, Writing-Original draft. **Tolulope Mafa-Attoye**: Conceptualization, Methodology, Data curation, Writing-Review and editing. **Megan Baskerville**: Data curation, Methodology, Investigation. **Eduardo Mitter**: Software, Visualization, Writing-Reviewing and editing. **Leandro Fonseca**: Methodology, Investigation. **Maren Oelbermann**: Conceptualization, methodology, Writing-Reviewing and editing. **Naresh Thevathasan**: Conceptualization, Funding acquisition, Project administration. **Siu Mui Tsai**: Conceptualization, Supervision. **Kari Dunfield**: Conceptualization, Project management, Supervision, Writing-Reviewing and editing.

## Competing Interests

The authors declare that they have no known competing financial interests or personal relationships that could have appeared to influence the work reported in this paper.

## Acknowledgements

We are thankful to Brian and Elizabeth Tew and Josie and Jens Madsen for providing access to their land during the study period. We thank Kira A. Borden by provide us with the data (laboratory analisis) on total carbon (TC) and nitrogen (TN). We are grateful to Kamini Khosla for technical support.

## Funding

This study was financially supported by research funds provided by the Agricultural Greenhouse Gas Program (AGGP) 12 administered by Agriculture and Agri-Food Canada, and the Natural Sciences and Engineering Research Council. Dasiel Obregon and Leandro Fonseca were supported by the São Paulo Research Foundation (FAPESP-2014/50320-4; 2015/13546-7; 2016/24695-6; 2018/09117-1; 2018/05223-1).

## Data availability

Most of the data have been included in the manuscript or the supplementary material. Data on DNA and cDNA sequencing will be made available on request.

## Notes

### Competing Interest Statement

The authors have declared no competing interest.

